# Ultrastructure expansion microscopy (U-ExM) of mouse and human kidneys for analysis of subcellular structures

**DOI:** 10.1101/2024.02.16.580708

**Authors:** Ewa Langner, Pongpratch Puapatanakul, Rachel Pudlowski, Dema Yaseen Alsabbagh, Jeffrey H. Miner, Amjad Horani, Susan K. Dutcher, Steven L. Brody, Jennifer T. Wang, Hani Y. Suleiman, Moe R. Mahjoub

## Abstract

Ultrastructure expansion microscopy (U-ExM) involves the physical magnification of specimens embedded in hydrogels, which allows for super-resolution imaging of subcellular structures using a conventional diffraction-limited microscope. Methods for expansion microscopy exist for several organisms, organs, and cell types, and used to analyze cellular organelles and substructures in nanoscale resolution. Here, we describe a simple step-by-step U-ExM protocol for the expansion, immunostaining, imaging, and analysis of cytoskeletal and organellar structures in kidney tissue. We detail the critical modified steps to optimize isotropic kidney tissue expansion, and preservation of the renal cell structures of interest. We demonstrate the utility of the approach using several markers of renal cell types, centrioles, cilia, the extracellular matrix, and other cytoskeletal elements. Finally, we show that the approach works well on mouse and human kidney samples that were preserved using different fixation and storage conditions. Overall, this protocol provides a simple and cost-effective approach to analyze both pre-clinical and clinical renal samples in high detail, using conventional lab supplies and standard widefield or confocal microscopy.

## Introduction

The development of expansion microscopy, which involves the physical expansion of biological samples for subsequent immunofluorescence imaging, has recently emerged as a valuable tool for both fundamental and clinical research applications. Expansion microscopy entails fixing the components of a specimen (e.g. a cell) to molecular anchors, crosslinkers that covalently connect target biomolecules (endogenous proteins, nucleic acids, or fluorescent probes bound to them), with a swellable acrylamide polymer. Then, the physical separation of the biomolecules is induced by swelling of the hydrogel with water. This allows for high-resolution structural analysis of macromolecular assemblies in cells and tissues with the use of regular diffraction-limited microscopes. Importantly, it eliminates the need for access to costly super-resolution microscopes to achieve sub diffraction-resolution fluorescence imaging of cellular structures. Although expansion microscopy was only recently established as a new imaging modality, several protocols have now been developed for expanding cells and tissues from a variety of organisms including bacteria, parasites, insects and vertebrates (Atchou et al., 2023; Bai et al., 2023; Bandeira et al., 2023; Campbell et al., 2021; Chang et al., 2024; Chang et al., 2017; Chen et al., 2015; Cheng et al., 2023; Ching et al., 2022; Chozinski et al., 2016; Chozinski et al., 2018; Damstra et al., 2022; Damstra et al., 2023; Dos Santos Pacheco & Soldati-Favre, 2021; Fan et al., 2021; Gallagher & Zhao, 2021; Gambarotto et al., 2021; Gambarotto et al., 2019; Gaudreau-Lapierre et al., 2021; Hamel & Guichard, 2021; Jurriens et al., 2021; Klimas et al., 2023; Kraft et al., 2023; Ku et al., 2016; Kunz et al., 2019; Kunz et al., 2021; Liffner et al., 2023; Lin et al., 2022; Louvel et al., 2023; Mäntylä et al., 2023; Middelhauve et al., 2023; Moye et al., 2023; Murakami et al., 2018; Park et al., 2019; Park et al., 2021; Parveen et al., 2023; Perelsman et al., 2022; Pernal et al., 2020; Pesce et al., 2019; Ponjavić et al., 2021; Rodriguez-Gatica et al., 2022; Sahabandu et al., 2019; Siegerist et al., 2022; Steib et al., 2022; Tillberg & Chen, 2019; Tillberg et al., 2016; Unnersjö-Jess et al., 2021; Unnersjö-Jess et al., 2018; Wainman, 2021; Wang et al., 2018; Wassie et al., 2019; Wen et al., 2023; Wilmerding et al., 2023; Woo et al., 2020; Yu et al., 2022; Zhao et al., 2017; Zhu et al., 2021; Zhuang & Shi, 2024).

Two main approaches have gained popularity over recent years, each with its own advantages and pitfalls. The initial protocol for expansion microscopy (ExM) developed by the Boyden laboratory (Chen et al., 2015) is based on expanding the immunofluorescent imprint of labeled factors after protein digestion. In this approach, the fluorescent moieties are first crosslinked to the gel polymer, followed by enzyme-based digestion of proteins, and subsequently expansion of the hydrogel (Chen et al., 2015; Chozinski et al., 2016; Tillberg et al., 2016; Zhang et al., 2016). This method requires very dense labeling (e.g. with antibodies) before the expansion step and is problematic for many proteins due to antibody-epitope hindrance after crosslinking. The second approach is termed the **m**agnified **a**nalysis of the **p**roteome (MAP) (Ku et al., 2016). MAP utilizes different crosslinking chemistry and, most importantly, involves protein denaturation with heat and detergent instead of enzymatic digestion. This has been shown to improve accessibility of otherwise blocked epitopes, resulting in better labeling and imaging. Several modifications of these two general methods have since been developed, including U-ExM (Gambarotto et al., 2021; Gambarotto et al., 2019), ExPath (Bucur & Zhao, 2018; Kylies et al., 2023; Zhao et al., 2017) and Magnify (Klimas et al., 2023), among others. These approaches are effective at expanding both normal and pathological cell/tissue samples, highlighting the potential utility of expansion microscopy as a research and diagnostic tool.

The detailed analysis of subcellular structures and organelles is still very challenging to perform in tissues, including the kidney. Even with the advent of super-resolution imaging systems, resolving ultrastructural changes in cytoskeletal elements in renal specimens is difficult, and the costly imaging systems are not widely available. Thus, the development of an imaging method that could be easily applied to kidney specimens, including pre-clinical and clinical samples, is crucial. In this study, we adapted and modified the MAP-based U-ExM protocol (Ching et al., 2022; Gambarotto et al., 2021; Gambarotto et al., 2019) to expand mouse and human kidney tissue sections (Figure 1). We describe a simple step-by-step protocol for the embedding, expansion, immunostaining, and imaging of diverse cytoskeletal structures. We highlight several markers of renal cell types, centrioles, cilia, the extracellular matrix, basement membrane, podocyte slit diaphragm, and several other cytoskeletal elements, all of which work well using this approach. Finally, we show that the method can be applied to both pre-clinical and clinical kidney samples, as well as different tissue fixation conditions. In sum, this protocol provides a cost-effective approach to analyze renal samples with near electron microscopy-level resolution, using existing lab supplies and standard fluorescence microscopy.

**Figure 1.**
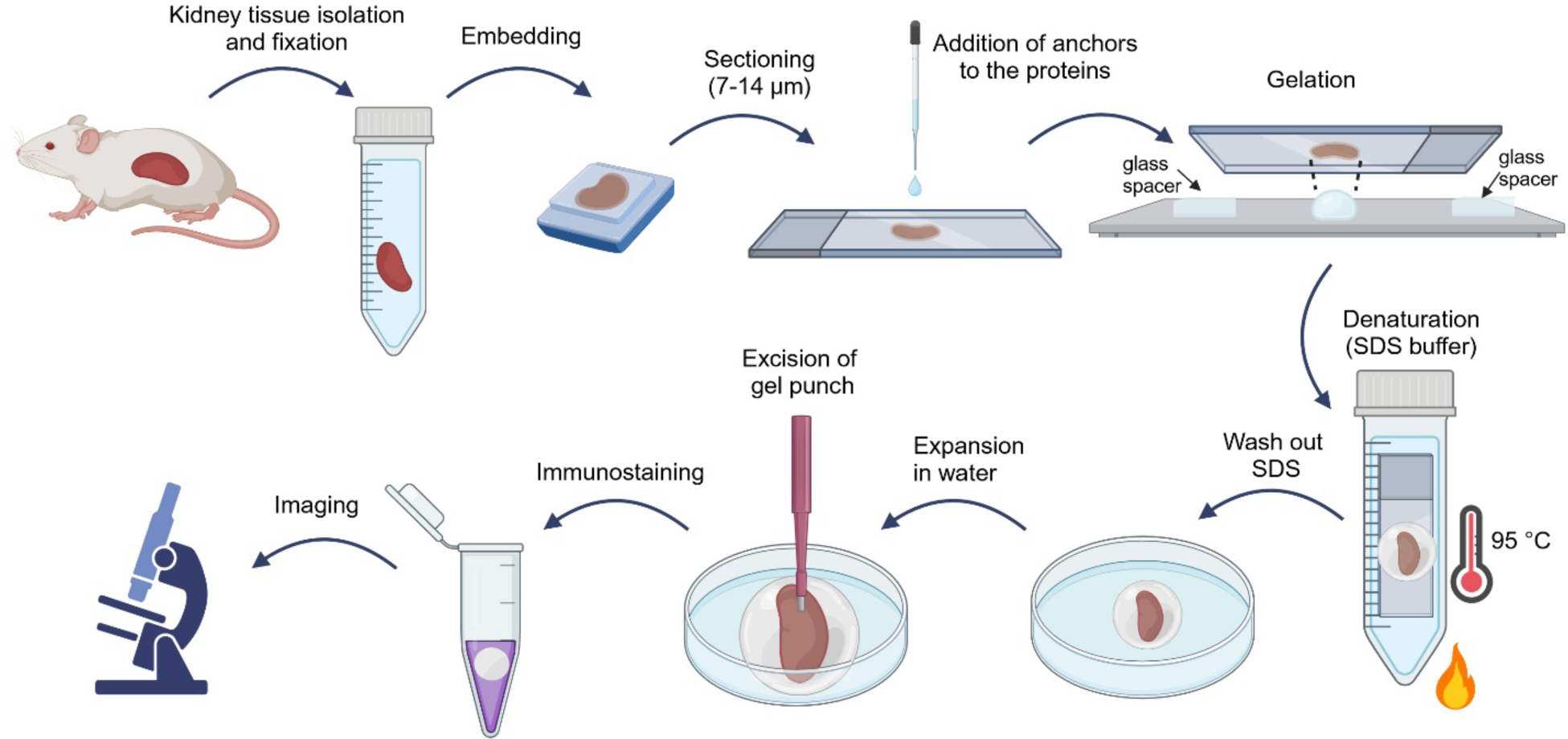
Overview of the kidney tissue section expansion steps using the modified U-ExM protocol.

## Materials and Methods

This protocol can be applied to pre-clinical (e.g. from mouse models) and clinical (human) kidney specimens, which were sectioned and placed onto microscope slides (Figure 1). For mouse, freshly isolated kidneys were fixed in 4% paraformaldehyde (PFA) overnight, then embedded in paraffin (stored at room temperature) or Optimal Cutting Temperature medium (OCT; kept frozen in -80 °C). For human kidney samples (obtained from the Kidney Translational Research Center, Washington University in St Louis), the tissue was fixed in either 4% PFA or 10% formalin, according to standard procedures. We observed successful expansion and antibody labeling with all the fixation/embedding conditions listed above. However, we discovered that the morphology of several structures was disrupted in non-fixed, freshly-frozen kidney tissue cryosections. Therefore, we recommend using traditionally well-fixed kidney specimens for expansion microscopy.

### 1. Tissue section preparation

#### PFA-fixed then OCT-embedded mouse kidneys

Kidneys are fixed in 4% PFA in PBS for 24 h, embedded in OCT and stored at -80 °C. We recommend preparing 10 μm thick kidney sections, placed on charged microscope slides and dried for up to 5 min at room temperature. Slides containing PFA-fixed/OCT-embedded kidney sections should be stored at -80 °C until ready for use. For these specimens remove the slide from the freezer, equilibrate at room temperature for 5 min, then proceed directly to Step 3 (“Addition of anchors to the proteins”). Removal (by peeling) of the OCT compound is optional and did not influence the subsequent expansion or labeling steps.

#### PFA-fixed then paraffin-embedded mouse and human kidneys

For mouse kidneys, the samples are fixed in 4% PFA in PBS for 24 h, dehydrated, embedded in paraffin (Hoshi et al., 2015) and stored at room temperature. For human kidneys, specimens may be either PFA (4%) or formalin (10%) fixed, and embedded in paraffin. We note that clinical human kidney samples fixed in buffered formalin tend to fragment during expansion, when compared with PFA-fixed tissue sections that retain their morphology better. Therefore, we recommend using PFA-fixed samples if accessible. Prepare 5-10 μm thick kidney sections, place them on charged microscope slides and bake for ∼16 h at 65°C to enable strong bonding of the tissue to the slide. Before moving to Step 3, the samples must be deparaffinized with xylene and rehydrated using decreasing (graded) ethanol washes, by following standard protocols. Following deparaffinization, the slides containing the kidney sections should be stored in water for 10 min before proceeding to Step 3.

### 2. Preparing solutions for expansion microscopy

(see Tables 1 and 2 for a complete list of reagents and materials required for the subsequent steps):

**Table 1:** Reagents needed.

| Reagent | Supplier (Catalog number) | Storage conditions |
| --- | --- | --- |
| Paraformaldehyde (PFA) 16% solution, EM grade | Electron Microscopy Sciences (15710 or similar) | Store at 4 °C |
| Embedding medium for frozen tissue specimens, Tissue-Tek® O.C.T. Compound | Sakura (4583 or similar) | Store at RT |
| Acrylamide (AA, 40%) | Sigma (A4058) | Store at 4 °C |
| N,N'-methylenebisacrylamide (BIS, 2%) | Sigma (M1533) | Store at 4 °C |
| Sodium Acrylate (SA, 97–99%) | Sigma (408220) | Store in desiccator at -20 °C |
| Formaldehyde (FA, 36.5–38%) | Sigma (F8775) | Store at RT |
| Ammonium persulfate (APS) | Bio-Rad (1610700) | Store in desiccator at RT |
| Tetramethylethylenediamine (TEMED) | Bio-Rad (1610800) | Store in RT |
| Poly-L-lysine solution, 0.1 % (w/v) in H<sub>2</sub>O – Ready to use | Sigma (P8920) | Store in RT |
| Sucrose, Methanol, SDS, NaCl, Tris, BSA, PBS (1x and 10x), Triton X-100, ddH<sub>2</sub>O water | Any supplier |  |

**Table 2.**
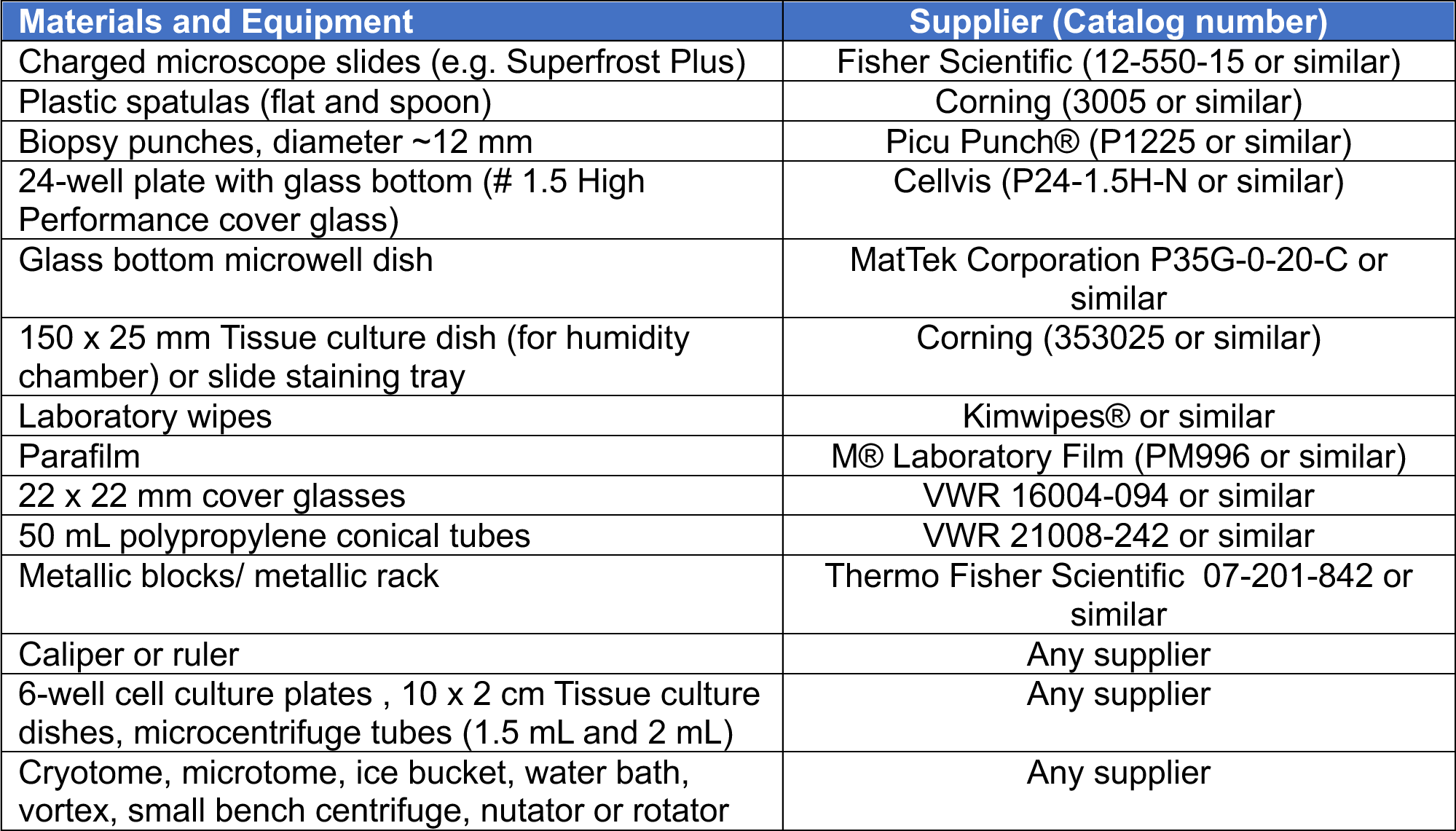
Materials and equipment.

#### Formaldehyde/acrylamide (0.7% FA / 1% AA) mix

Prepare this solution fresh immediately before use. Combine 19 μL of FA and 25 μL of AA (see Table 3) with 956 μL of 1× PBS, for a final concentration of 0.7% FA and 1% AA (this yields 1 mL of 1× solution). This volume is sufficient for at least 6 slides/specimens, with an area of ∼100 mm^2^ marked with a hydrophobic slide marker (PAP-pen; see Step 3). If a larger tissue sample is used, increase the volume of the FA/AA mix according to the surface area of the sample.

**Table 3.** Formaldehyde/acrylamide mix solution.

| Reagent | Stock concentration | Working concentration | Volume (μl) |
| --- | --- | --- | --- |
| Formaldehyde | 36.5-38% | 0.7% | 19 |
| Acrylamide | 40% | 1% | 25 |
| PBS | 1x |  | 956 |

#### Sodium acrylate (SA) stock solution

Prepare 38% (w/v) stock solution by dissolving 3.8 g of SA in 10 mL final volume of autoclaved ddH_2_O water. Once the SA fully dissolves, keep the solution at 4 °C *(Caution: Work in a fume hood, wear gloves and mask).* Note that at this step it is crucial to ensure high quality of SA solution. Remember to use only clear or slightly yellowish (but not strong yellow) SA solutions to avoid problems with gel polymerization and expansion. Discard solutions with strong yellow color or containing precipitate. Since the quality and purity of SA may vary depending on the lot, test every new batch before using in your experiments. Good quality SA solutions may be stored at 4 °C until it turns yellow.

#### U-ExM monomer solution (MS; 19% SA/ 10% AA/ 0.1% BIS-AA in 10x PBS)

Table 4 indicates the volumes of SA, AA and Bis-AA used to prepare 450 μL of monomer solution (MS). Mix all the reagents thoroughly, make aliquots of 50 μL per reaction and keep on ice until next step *(Caution: Work in a fume hood and wear gloves)*.

**Table 4.** Monomer solution.

| Reagent | Stock concentration | Working concentration* | Volume (μl) |
| --- | --- | --- | --- |
| Sodium acrylate (SA) | 38% | 19% | 250 |
| Acrylamide (AA) | 40% | 10% | 125 |
| Bis-Acrylamide (BIS-AA) | 2% | 0.1% | 25 |
| PBS | 10x | 1x | 50 |
| Final volume |  |  | 450 |

\* The following values correspond to the final concentrations in the gelling solution after TEMED and APS are added (see section 2).

#### Solution for gelation (MS with 0.55% TEMED and APS)

Prepare 10% stock solutions of TEMED (free radical accelerator) and APS (free radical initiator). For 10 μL of 10% TEMED, dilute 1 μL of the main stock (100%) with 9 μL of ddH_2_O water. For 10% APS, dissolve 50 mg in ddH_2_O to a final volume of 500 μL. Make aliquots fresh before each experiment. To prepare geling solution, take an aliquot of 50 μL of MS and add 3.12 μL of TEMED then 3.12 μL of APS from the 10% stocks immediately before the polymerization step (Step 3, “Gel polymerization”). This brings the final concentrations of TEMED and APS to 0.55%.

#### Denaturation buffer (200 mM SDS/200 mM NaCl, 50 mM Tris, pH9)

Table 5 indicates volumes and weights used for the preparation of 1000 mL of denaturation buffer. Start by dissolving 6.06 g of Tris-Base in 500 mL of ddH2O in a glass beaker. Add 11.69 g of NaCl and 57.68 g of SDS while stirring. Adjust to pH9 with HCl and bring to 1000 mL with ddH2O.

**Table 5.** Denaturation buffer.

| Reagent | Amount (g) | Working concentration |
| --- | --- | --- |
| SDS | 57.68 | 200mM |
| NaCl | 11.69 | 200mM |
| Tris-Base | 6.06 | 50mM |
| ddH<sub>2</sub>O | Fill up to 1000 mL |  |

### 3. Expansion microscopy procedure

#### Day 1

##### Addition of anchors to the proteins

Before proceeding to the next step, prepare the slides containing kidney tissue appropriately depending on preservation type (described in Step 1).

a. Draw a circle around the tissue section with a hydrophobic slide marker (Pap-Pen). We usually mark an area of ∼100 mm^2^ (use a 12 mm round coverslip as a guide if it helps).
b. Prepare the gelation humidity chamber (Figure 2A). Add a piece of parafilm where the slide will be deposited.
c. Place slides in the humidity chamber, add 100 μL of 1× PBS and incubate for 5 min. Aspirate and repeat this wash step with PBS twice more.
d. Remove PBS and add 150 μL of AA/FA mix to each section, then incubate for 2 h at room temperature in the humidity chamber.
e. In the meantime, place the metallic blocks and a metal tube rack (see Table 2) in the -20 °C freezer to ensure they are fully chilled.

**Figure 2.**
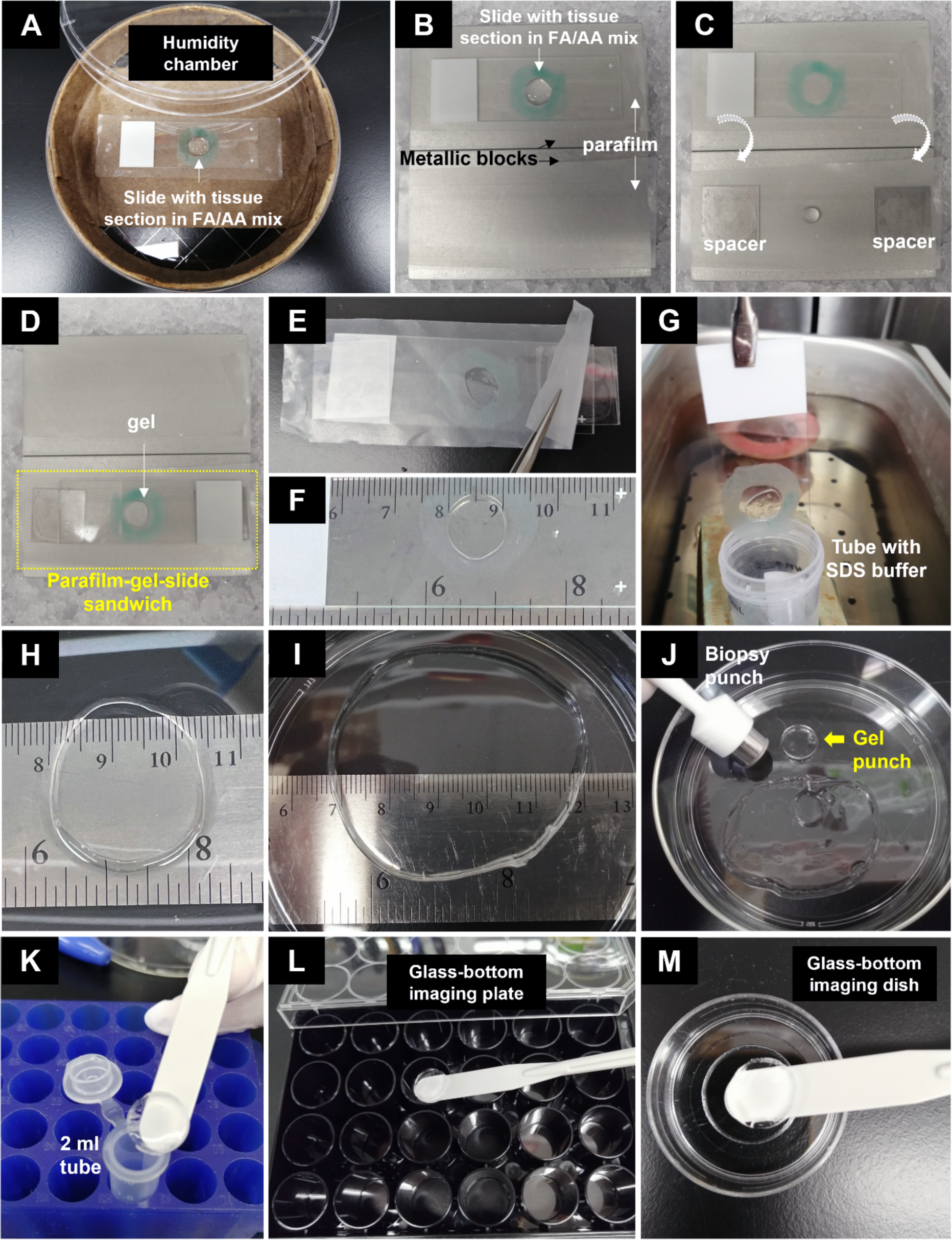
Steps of the U-ExM protocol for kidney tissue sections. **(A)** Addition of anchors to the proteins in the tissue section. **(B)** Assembly of the gelation station. **(C-D)** Slide preparation and gel polymerization. **(E-F)** Measuring dimensions of the gel before expansion. **(G)** Protein denaturation in SDS buffer with heat. **(H)** Partial gel expansion post-denaturation. **(I)** Final gel size after expansion in water. **(J)** Obtain gel pieces using a biopsy punch. **(K)** Transfer individual gel punch into tube for immunostaining. **(L-M)** Imaging of gels. Transfer stained and expanded gel punch (after removing excess water with a Kimwipe) into a glass-bottom imaging plate or round dish. Remove as much water as possible before transfer to avoid gel movement during imaging.

##### Gel polymerization

(*Note: To ensure proper gelation this step must be performed on ice keeping all the reagents cold. Low temperature slows the polymerization rate, an essential factor for gelation. Prepare the following solutions immediately before starting the gelation steps):*

a. Determine the total volume of monomer solution (MS) needed. Each sample/slide requires 100 μL of solution, which includes enough MS for the wash step (we recommend preparing an extra 100 μL to account for loss during pipetting).
b. Prepare 10% TEMED and APS. You will require 3.12 μL of each per slide. Keep cold on ice.
c. In order to be as fast as possible, we advise using two pipettes (e.g. a P10 set at 3.12 μL for TEMED and APS, and a P100/200 set at 50 μL for the MS solution).
d. Move the metallic tube rack and metallic blocks from -20 °C to a bucket filled with ice (Figure 2B). Place a piece of parafilm directly on the two metallic blocks where the slides will be deposited.
e. Prepare the MS (make enough total volume for all slides in one 1.5 mL tube). Aliquot 50 μL of MS into separate 1.5 mL tubes (one for each slide), and keep them chilled in the metallic rack on ice.
f. Take out a slide from the humidity chamber and place on one of the metallic blocks covered with parafilm (Figure 2B). The slide should still contain the FA/AA mix. We recommend proceeding with one slide at a time.
g. On the second block covered with parafilm place two “spacers” (each spacer is made by stacking three 22×22 mm coverslips). Position the spacers so that the edges of the glass slide can easily sit on them (Figure 2C).
h. Aspirate the FA/AA mix from the slide, add 50 μL MS solution <u>without</u> TEMED or APS to the tissue section, and incubate for 1-2 min.
i. <u>Fast steps</u>: Add 3.12 μL of TEMED and 3.12 μL of APS to one of the 1.5 mL tubes containing 50 μL aliquot of MS, pipette 4-5 times while keeping the tubes on ice. To be even faster, you do not need to close the tube lid during sample mixing. Note that TEMED should be added first to delay free radical formation for as long as possible. Because of the presence of TEMED and APS, gelation will start in a few seconds, so these (and subsequent steps) must be performed extremely rapidly.
j. Place 50 μL of the activated MS on the parafilm between the spacers (Figure 2C). This volume may be increased for larger tissue sections.
k. Quickly remove the inactive MS solution covering the sample and flip the slide placing the tissue section directly onto the 50 μL drop of activated MS (Figure 2C-D).
l. Press both ends of the slide (i.e. the regions sitting on the spacers) gently with your fingers, keep pressing and holding it in place for 2 min as the gel slowly polymerizes, then leave it on ice for an additional 5 min to let the MS penetrate the tissue while polymerization occurs.
m. Ensure that the gel has become visibly solid, then transfer the parafilm-gel-slide assembly to a humidity chamber (keep it in the same orientation), and incubate overnight at room temperature.

#### Day 2

##### Protein denaturation

a. Remove the parafilm-gel-slide from the humidity chamber. Flip the sandwich (parafilm side facing up) and gently peel the parafilm away from the gel (Figure 2E). The gel should remain tightly stuck to the slide.
b. Using a ruler (or caliper), measure the dimensions of the gel before denaturation (Figure 2F). *Note: It is crucial to measure the gel size at this stage before proceeding with the denaturation step, as the gels will slightly expand during this process. This value represents the “pre-expanded” gel size*.
c. Transfer the slide with the gel into a 50 mL conical tube filled with 35 mL of denaturation buffer (use one slide per tube; Figure 2G).
d. Place the tubes in a pre-heated 95 °C water bath (alternatively, you can use a beaker of water heated to 95 °C on a hotplate). Incubate the gel-containing slide for 4 h at 95 °C.
e. Remove the conical tube from the water bath and let it cool down at room temperature (∼30 min) before proceeding to the next step. *Not*e: *The gel will completely detach from the slide during the 95 °C incubation step and will be found at the bottom of the tube. The gel will have slightly expanded at this stage, which is to be expected (Figure 2H)*.

##### Gel expansion

a. Remove the glass slide from the 50 mL conical tube and discard, then slowly pour out the denaturation buffer making sure that the gel does not fall out.
b. Gently transfer the gel into a 100 mm tissue culture dish filled with 50 mL of ddH_2_O water. Place only one gel per dish, as each one will expand to form a large gel (Fig. 1H). Incubate for 20 min at room temperature.
c. Exchange the water with fresh 50 mL of ddH_2_O, being careful not to disrupt the gel’s integrity, and incubate for another 20 min. Repeat this wash step three more times. *Note: as the gel starts to expand it may slightly roll up around the edges. This is normal, and will resolve itself during the subsequent incubation step*.
d. After the final wash, incubate the gel in 50 mL of ddH_2_O overnight at room temperature.

#### Day 3

Discard the water and measure the gel dimensions (Figure 2I). This will help to determine the expansion coefficient at the macro scale. *Note: After the overnight incubation the gel should expand to ∼4 times their initial size (compared to its size after gelation, Figure 2F).* Store the gel in ddH_2_O at 4 °C until needed for immunostaining. The gel will remain stable and can be stored in water for at least ∼4 months in a sealed culture dish to avoid evaporation.

##### Immunostaining

a. Cut out small (12 mm diameter) gel pieces using a biopsy punch (Figure 2J). *Note: the tissue will generally be in the center of the gel, so start with acquiring punches form the central region*.
b. Transfer the punched-out piece of gel (“gel punch”) to a 2 mL tube containing 1.8 mL PBS.
c. Wash the gel punch gently at least two times (15 min each) in PBS on a rotator/nutator at room temperature. *Note: During this step, the gel will decrease to about half of its expanded size due to water loss. This is performed to avoid diluting of antibodies in the following steps and allow for smaller volumes of antibody solutions to be used. Be sure that you do not accidently aspirate or puncture/break the gel with a pipette tip during wash steps*.
d. Remove the PBS and replace with 200 μL of primary antibodies diluted in PBS-BT (PBS containing 3% BSA and 0.1% Triton X-100). *Make sure that the gel punch is placed at the bottom of the tube and antibody solution is covering it*.
e. Incubate at 4 °C overnight (preferrable) or at 37 °C for 2-3 h.

The full list of antibodies used in this study and shown to be effective is provided in Table 6. We recommend starting with an antibody concentration that is 2× that used for standard immunofluorescence. Note that some antibodies may perform better when incubated at 37 °C, or for shorter periods of time. Incubation times will need to be empirically determined.

**Table 6.** Antibodies used in this study.

| Antibodies (species) | Source | Cat. No. |
| --- | --- | --- |
| Acetylated α-tubulin (Mouse IgG2b) | Sigma | T6793 |
| Actin (Rabbit) | Cell Signaling | 4968 |
| Aquaporin 1 (Rabbit) | Abcam | 168387 |
| Aquaporin 2 (Mouse IgG1) | Santa Cruz Biotechnology | sc-515770 |
| Arl13b (Rabbit) | Proteintech | 17711-1-AP |
| Centrin (Mouse IgG2a) | Millipore | 04-1624 |
| Cep120 (Rat) | Betleja et al. (2018) | Homemade |
| Cep135 (Rabbit) | Abcam | ab75005 |
| CLC-K (Rabbit) | Alomone Lab | ACL 004 |
| Collagen IV (Goat) | Southern Biotech | 1340-01 |
| Desmin (Mouse IgG1) | Agilent Technologies | M0760 |
| E-cadherin (Mouse IgG2a) | BD Bioscience | BDB610181 |
| Glutamylated tubulin (GT335; Mouse IgG1) | AdipoGen | AG20B-0020-C100 |
| Nephrin (Goat) | R&D Systems | AF3159 |
| Odf2 (Rabbit) | Sigma | HPA001874 |
| Podocalyxin (Goat) | R&D Systems | AF1554 |
| Podocin (Rabbit) | IBL International | JP29040 |
| Rootletin (Rat) | Kind gift from Didier Hodzic | Homemade |
| Synaptopodin (Guinea Pig) | Synaptic Systems | 163004 |
| Synaptopodin (Guinea Pig) | American Research Products | 03-GP94-N |
| α-Actinin 4 (Rabbit) | ThermoFisher Scientific | 42-1400 |
| γ-tubulin (GTU-88; Mouse IgG1) | Sigma-Aldrich | T6557 |

#### Day 4

##### Secondary antibody labeling

a. Wash the gel punch with 1.8 mL of PBST (1× PBS supplemented with 0.1% TritonX-100) three times (10 min each) at room temperature on a nutator.
b. Remove the final wash solution and add 200 μL of secondary antibodies diluted in PBS-BT (1× PBS supplemented with 0.1% TritonX-100? and 3% BSA).
c. Incubate for 3 h at 37 °C on a rotator/nutator and protect from light. *Note: Use of a nutator during incubation with antibodies is optional. If needed, you can also perform secondary antibody staining overnight at 4 °C. For nuclear DNA staining with DAPI, incubation overnight at 4 °C is required*.
d. Wash the gel three times with PBST (10 min each) at RT on a nutator.

Final gel expansion prior to imaging:

Transfer the punched gel piece into one well of a 6-well plate containing 5 mL of ddH_2_O. Place only one gel punch per well. Incubate for 30 min to remove all traces of salt from the final PBST wash. Exchange with fresh 5 mL of water and incubate for another 30 min. *Note: Remember to protect the samples from light during these steps*.

##### Gel imaging

We recommend using an inverted microscope with the gel punch placed in either a glass-bottom 24-well plate or a 35 mm imaging dish (see Table 2).

a. Using a flat plastic spatula, carefully transfer the gel punch onto a Kimwipe to remove residual water from the gel.
b. Transfer the gel into the well of an imaging plate (Figure 2L-M). (*Note: at this step you will not be able to easily identify which side of the gel contains the tissue. The tissue-containing side of the gel will appear to be slightly less “shiny”, which may help in identifying the correct orientation. However, if you place one side of the gel down on the glass and are unable to locate the cells on the microscope, you will need to flip the gel over to see if the samples are present on the other side. We recommend starting with a low magnification objective lens to first locate the side of the gel containing the tissue, before moving to higher magnification objectives)*.
c. Gently push down on the gel to remove any water or air gap between the gel and the glass. (*Note: to help reduce gel sliding during imaging, the glass-bottom imaging dishes can be pre-coated with poly-L-lysine. However, this may result in the gel sticking to the dish. Therefore, be sure to capture images of all needed fields of your specimen since you might not be able to reuse the gel again)*.
d. In the following experiments, we used a Nikon Eclipse Ti-E inverted confocal microscope equipped with a 20x (0.75NA), 40x (1.3NA), 60× (1.4NA) and 100x (1.45NA) Plan Fluor oil immersion objective lens (Nikon, Melville, NY). A series of digital optical sections (Z-stacks) were captured at 0.2 - 0.3 µm intervals using a Hamamatsu ORCA-Fusion Digital CMOS camera. Image volumes were deconvolved using Nikon Elements software. For improved resolution, we also captured images on a Zeiss LSM880 Airyscan confocal microscope equipped with a 40x (1.3NA) and 63× (1.4NA) oil immersion objective. Z-stack images were obtained at 0.16 to 0.30 µm intervals and were analyzed using ImageJ2 or Volocity (Quorum Technologies Inc.) software.

Following imaging, the gel samples can be stored in water at 4 °C. We highly recommend performing imaging within 2 weeks post-staining, to ensure that the quality of the gel, and the fluorescence signals, are maintained.

## Results

### Analysis of centrioles and cilia

First, we sought to determine the expansion factor of the kidney samples following U-ExM, by using the well-defined size of centrioles as a measuring tool. Centrioles have a conserved and defined size across different eukaryotic cells: they are ∼250 nm in diameter and ∼500 nm in length (Avidor-Reiss & Gopalakrishnan, 2013; LeGuennec et al., 2021). Therefore, they can serve as an internal subcellular ruler. We used a robust marker of centriolar microtubules to visualize and measure their dimensions *in situ* after expansion. Gels were immunostained with antibodies against acetylated α-tubulin to label cilia and centrioles, as well as E-cadherin to mark cell junctions (Figure 3A-C). We observed strong labeling of cilia and centrioles in nephron tubular epithelial cells in the expanded tissues (Figure 3B-F), consistent with that seen in unexpanded samples. Thus, acetylated α-tubulin is also a reliable marker for analysis of centrioles and cilia morphology in expanded kidney tissues.

**Figure 3.**
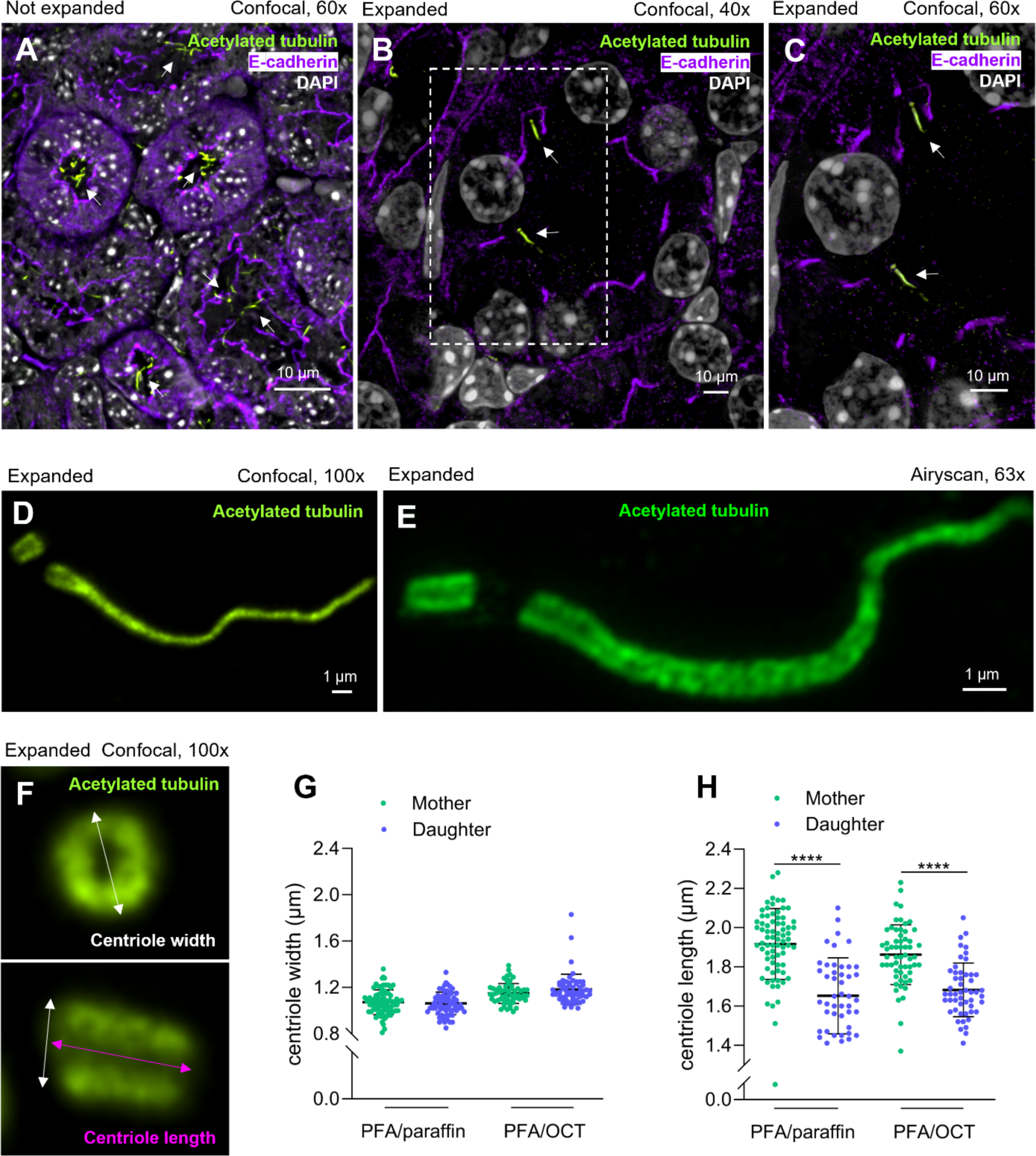
Analysis of centrioles and cilia in mouse kidney tubular epithelial cells. **(A-C)** Immunofluorescent labeling of the centrosome-cilium complex in regular (A) versus expanded (B-C) kidney sections of P15 mouse kidneys. Centriolar and ciliary microtubules were labeled with antibodies against acetylated α-tubulin (green), nephron tubular epithelia with E-cadherin (purple), and nuclei with DAPI (white). Arrows point to cilia in nephron tubules. **(D-E)** High-magnification images of centrioles and cilia in expanded kidney sections imaged by (D) confocal or (E) Superresolution microscopy. **(F)** Representative images used for quantification of centriole size. White: outer centriole diameter; Magenta: centriole length. **(G)** Quantification of mother and daughter centriole width in expanded PFA/paraffin (n=147) and PFA/OCT (n=117) preserved kidney sections. **(H)** Quantification of mother and daughter centriole length in expanded PFA/paraffin (n=121) and PFA/OCT (n=110) preserved kidney sections. Data were analyzed using two-tailed unpaired *t* test. The vertical segments in box plots show the first quartile, median, and third quartile. The whiskers on both ends represent the maximum and minimum values for each dataset analyzed. **** p< 0.0001. Experiments were not blinded.

To quantify the expansion factor in our samples at the nanometer scale, we measured the dimensions (width and length) of centrioles that serve as the gold standard for various protocols of ExM (Ching et al., 2022; Gambarotto et al., 2021; Gambarotto et al., 2019; Louvel et al., 2023; Sahabandu et al., 2019). Centriole width and length were measured in expanded gels from both paraffin and OCT-embedded kidney sections (Figure 3F-H). In PFA-fixed/paraffin-embedded kidneys, the mean diameter of both mother and daughter centrioles was 1.07 μm and 1.06 μm, yielding an expansion factor of 4.3× and 4.2×, respectively (calculated based on using 250nm as the expected width of a centriole). The mean length of the mother centriole was 1.91 μm, indicating an expansion coefficient of 3.8x (assuming a length of 500nm). The daughter centriole was slightly shorter (Figure 3F-H), as expected. In PFA-fixed/OCT-embedded kidneys, the mean diameter of both mother and daughter centrioles was 1.14 μm and 1.18 μm (expansion factor of 4.6× and 4.7×; Figure 3F,G). The length of the mother centriole was 1.86 μm (expansion factor of 3.7×; Figure 3F,H). Therefore, the two fixation conditions yield similar results. These values are consistent with the expansion factor of the hydrogel calculated at the macro scale (∼4x; Figure 2 and Step I). Moreover, they are consistent with results from other laboratories using expansion microscopy protocols on cells grown in culture (Bucur & Zhao, 2018; Campbell et al., 2021; Chang et al., 2017; Chen et al., 2015; Chozinski et al., 2016; Chozinski et al., 2018; Gambarotto et al., 2021; Gambarotto et al., 2019; Sahabandu et al., 2019; Zhang et al., 2016; Zhu et al., 2021). Altogether, our results indicate that applying this U-ExM protocol to kidney samples will provide isotropic expansion of the tissue, which can be verified at the nanometer scale.

### U-ExM is effective for analysis of centrosomal and ciliary protein composition

Next, we tested whether U-ExM can be used to finely localize other centriolar and ciliary proteins in expanded mouse kidney tissue. We tested several commercially available antibodies that are commonly used in the field to label various components of ciliary and centrosomal apparatus. This includes ciliary axoneme and membrane markers, pericentriolar material, centrosome rootlets, as well as centriolar lumen, cartwheel, and appendages proteins. Of note, these substructures are often disrupted in patients with renal ciliopathies (Devlin & Sayer, 2019; Fleming et al., 2017; McConnachie et al., 2021; Reiter & Leroux, 2017; Srivastava et al., 2018), making the ability to visualize them in nanoscale resolution important for basic and clinical research. As ciliary markers, we found that U-ExM is useful for detecting glutamylated tubulin (Figure 4A; marks both centriolar and ciliary microtubules) (Wloga et al., 2017), and Arl13b (Figure 4B) (Larkins et al., 2011). Several markers of centriole substructures were also robust: Cep120 (Figure 4C) was localized to both centrioles but enriched on the daughter centriole (Betleja et al., 2018; Mahjoub et al., 2010); the cartwheel assembly protein Cep135 (Figure 4D) was present on both centrioles, with higher expression evident on the mother centriole (Dahl et al., 2015); the inner core protein Centrin (Figure 4E) was localized in the distal lumen of both centrioles (Salisbury, 1995); the subdistal appendage component Odf2 (Figure 4F) was localized only on the mother centriole in the expected region of the subdistal appendages (Ishikawa et al., 2005); the pericentriolar material component γ-Tubulin (Figure 4G) was detected as a “cloud” surrounding the two centrioles of the centrosome (Khodjakov & Rieder, 1999); and the centriolar rootlet factor Rootletin (Figure 4H) was localized to inter-centriole linker fibers (Vlijm et al., 2018; Yang et al., 2002). Therefore, we found that the U-ExM protocol is useful for studying the composition of multiple substructures of the centrosome-cilium complex in remarkable detail using expanded kidney tissue sections.

**Figure 4.**
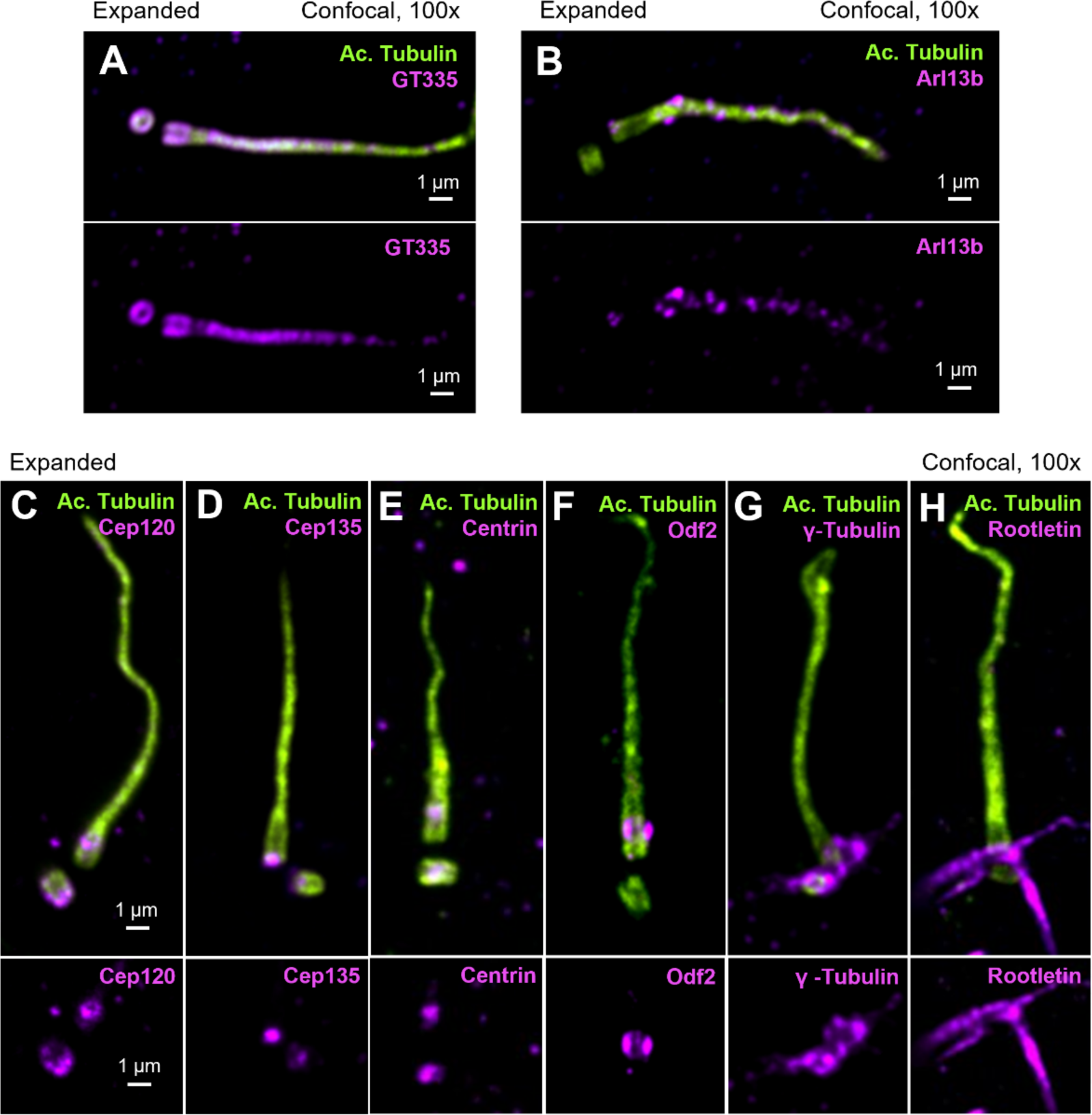
Validation of ciliary and centriolar markers using U-ExM of mouse kidneys. Representative immunofluorescence images of expanded P15 mouse kidneys stained with antibodies to mark **(A)** the ciliary axoneme and centriolar microtubules with acetylated (green) and polyglutamylated (magenta) α-tubulin, **(B)** ciliary GTPase Arl13b (magenta), **(C)** centriole assembly factor Cep120, **(D)** centriolar cartwheel component Cep135**, (E)** centriole apical lumen protein Centrin, **(F)** subdistal appendage protein Odf2, **(G)** the pericentriolar material marker γ-tubulin, and **(H)** the ciliary rootlets component Rootletin (all in magenta).

### Validation of kidney cell type-specific markers

Next, we explored whether U-ExM can be used to label and distinguish between specific compartments and cell types of the kidney. This question is important for both fundamental and clinical research applications, since different renal diseases will involve defects in only certain populations of cells, making it essential to identify the cell types of interest by microscopy. We tested several commercially available antibodies that specifically mark the various segments of nephron tubules, as well as collecting ducts, glomerular, and stromal cell markers. We observed robust labeling of the proximal tubule (Figure 5A) using Aquaporin 1 (Verkman, 2002), distal tubule (Figure 5B) using CLC-K (Kieferle et al., 1994), and the collecting duct (Figure 5C) using Aquaporin 2 (Verkman, 2002). There was concomitant labeling of all epithelial cells along the nephron tubule with E-cadherin (Figure 5A) (Prozialeck et al., 2004). Collagen IV (Figure 5B) marked the basement membranes (Miner, 1999), and actin staining was robust in the majority of tubular and stromal cells (Figure 5C). Finally, we examined markers of different cell types found in the kidney glomerulus, such as podocytes (Synaptopodin) and mesangial cells (Desmin), both of which clearly labeled the appropriate structures (Figure 5D) (Mundel et al., 1997; Stamenkovic et al., 1986). Collectively, we conclude that U-ExM of kidney sections allows for labeling and identification of specific cell types of interest for further analyses.

**Figure 5.**
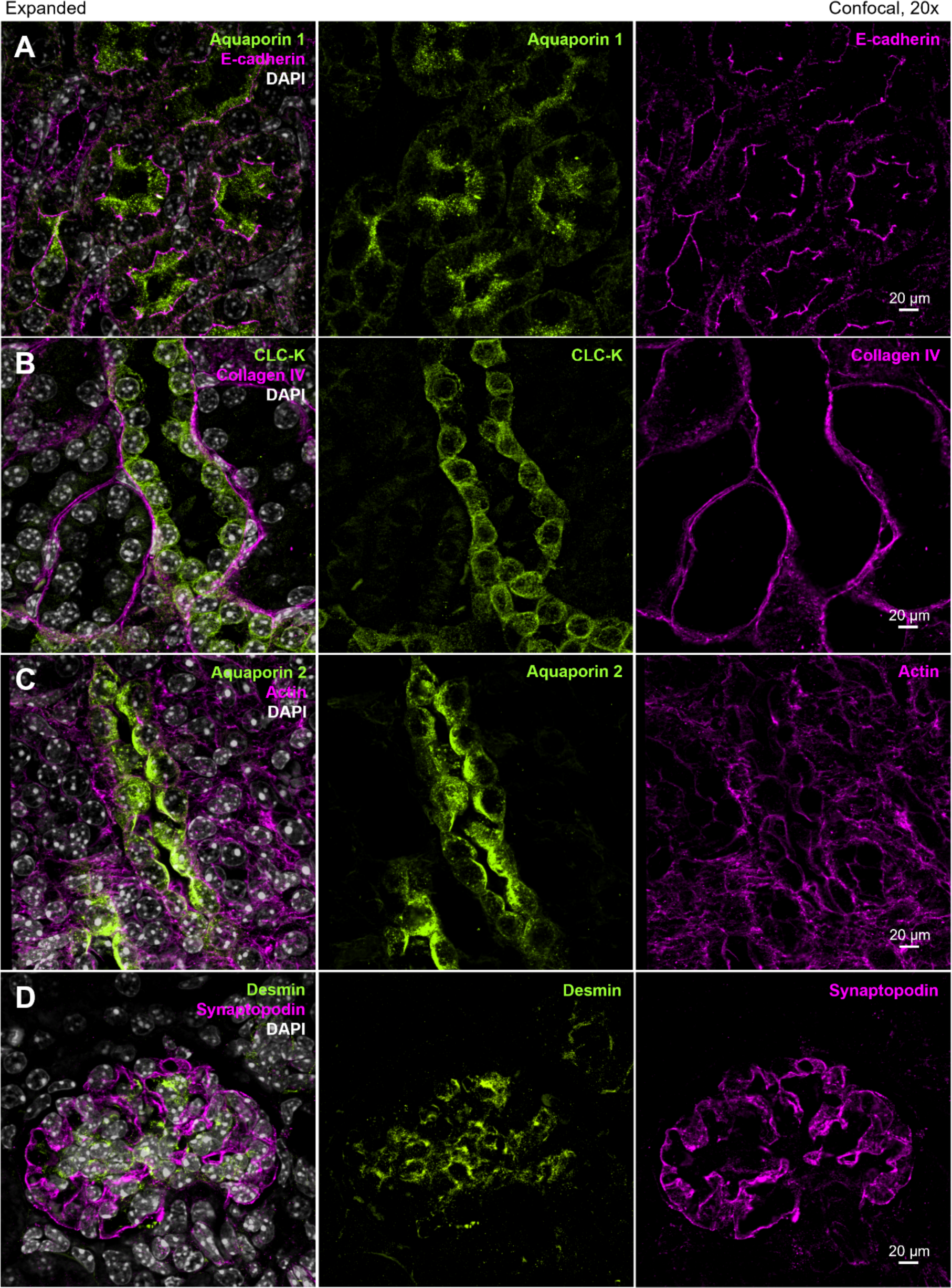
Testing of nephron segment and cell type-specific kidney markers. Example immunofluorescence images of expanded P15 mouse kidney sections stained with antibodies to mark **(A)** proximal tubule epithelial cells with Aquaporin 1 (green) and epithelial cell junctions with E-cadherin (magenta), **(B)** distal tubules with CLC-K (green) and the basement membrane with Collagen IV (magenta), **(C)** collecting ducts with Aquaporin 2 (green) and the actin cytoskeleton (magenta), **(D)** glomerular podocytes with Synaptopodin (magenta) and mesangial cells with Desmin (green). Nuclei were counter-stained with DAPI (white).

### Analysis of glomeruli and the podocyte slit diaphragm

A major challenge in the nephrology field is the distinction between healthy podocyte foot process and foot process effacement, a morphological change that indicates podocyte injury (Kriz et al., 2013). These studies have been dependent on electron microscopy-based analysis, which is essential to visualize the changes in these nanoscale foot processes, and the slit diaphragm that bridge the gap in between the adjacent foot processes. To determine whether U-ExM would be effective at resolving these structures using fluorescence microscopy, we expanded then immunostained mouse kidney sections (normal, non-diseased) with antibodies against the extracellular domain of Nephrin, a major slit diaphragm protein (Ruotsalainen et al., 1999; Wartiovaara et al., 2004), and Synaptopodin, a podocyte-specific actin-associated protein (Mundel et al., 1997). Imaging of the expanded samples clearly defined the podocyte foot process boundaries, with the slit diaphragm located in between them (Figure 6A and B). In cross-sectional views, Synaptopodin clusters representing the individual foot processes were flanked by Nephrin staining in between, in positions that are consistent with the locations of the slit diaphragms (Figure 6C). Co-staining with two slit diaphragm proteins, Nephrin and Podocin (Roselli et al., 2002), showed that Nephrin is present in the center of the slit diaphragm, with Podocin on the interdigitating foot processes, forming two leaflets on both sides, representing cell membrane staining (Figure 6D-H).

**Figure 6.**
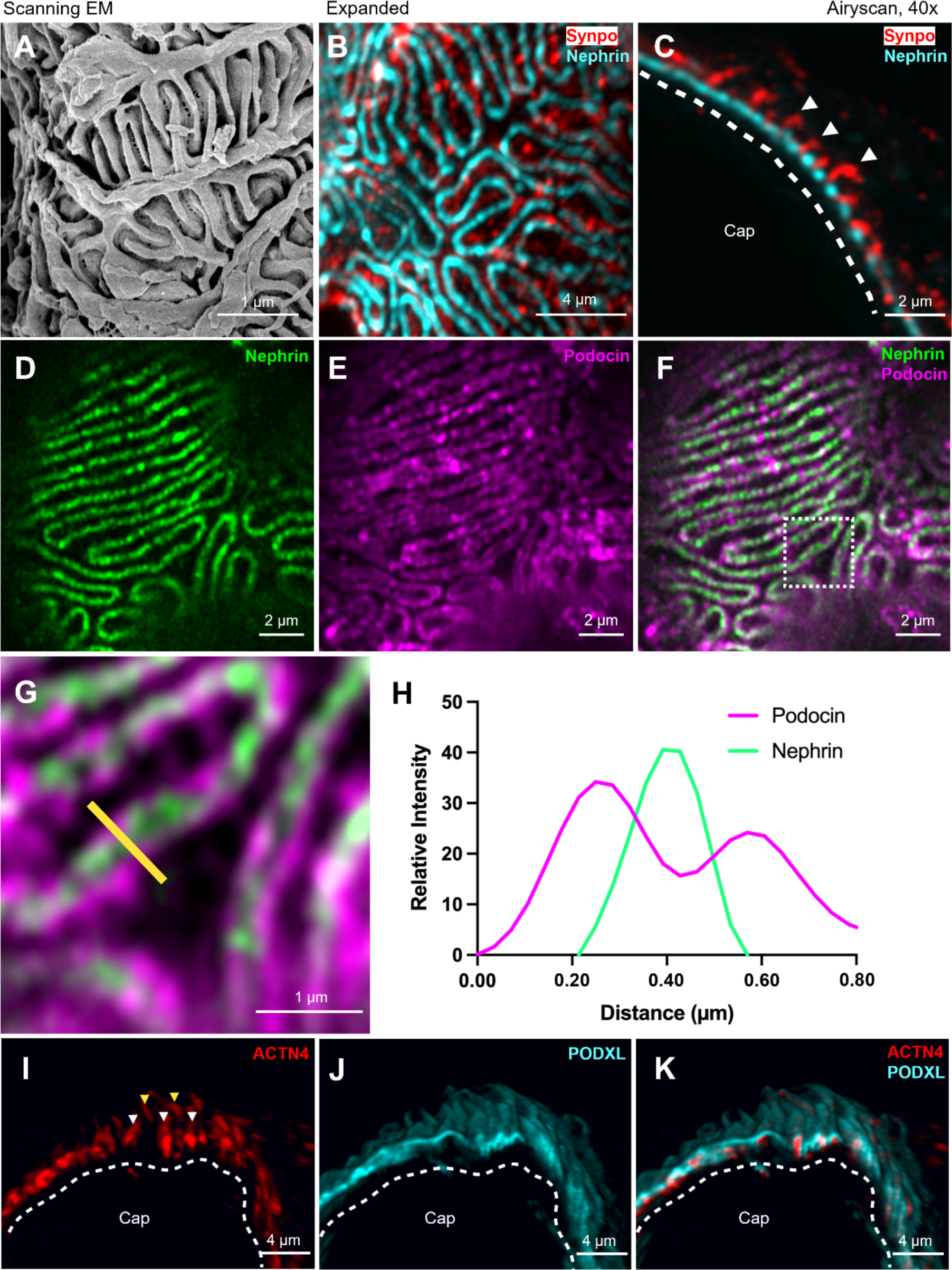
Analysis of expanded glomerular podocytes. **(A)** Scanning electron microscope (SEM) image of a mouse kidney highlighting the interdigitating foot processes and slit diaphragms. **(B-C)** Expanded P40 mouse kidney sections labeled for Nephrin (cyan) and Synaptopodin (Synpo, red), showing localizations of the slit diaphragm and podocyte foot process, respectively (*en face:* B; cross-sectional views: C). Arrowheads highlight individual foot processes and the dashed line denotes the expected location of the glomerular basement membrane. **(D-G)** Example *en face* images of two slit diaphragm markers, Nephrin (green) and Podocin (magenta). **(H)** Fluorescence intensity map analysis of the slit diaphragm displays a Nephrin peak signal in the center (green), flanked by two Podocin peaks from adjacent foot processes. The Plot Profile function on ImageJ was used to create an intensity plot along a straight line drawn perpendicularly. **(I-K)** 3D reconstruction of a glomerular capillary wall (Cap) labeled with antibodies against the actin-bundling protein α-actinin 4 (Acnt4, red) and Podocalyxin (Podxl, cyan). White arrowheads denote foot processes from an adjacent podocyte indicated by yellow arrowheads.

Finally, we performed 3D reconstruction of the samples co-labeled with Podocalyxin (Podxl, a transmembrane sialoglycoprotein) (Kerjaschki et al., 1984), together with α-actinin 4 (ACTN4, the major actin-bundling protein in the podocytes) (Lachapelle & Bendayan, 1991). We resolved the interdigitating foot processes outlined by Podxl as it stains the lateral and apical membranes of individual podocyte foot processes (Figure 6J, K). This signal was surrounding the ACTN4 staining that labels the foot processes in a linear fashion along the longitudinal axis (Figure 6I). Altogether, these experiments highlight the feasibility of resolving foot processes and slit diaphragms of podocytes, provide further quantitative proof that the U-ExM of the kidney tissue is occurring at the micro scale, and that the technique will be a useful to study pathologies associated with podocytes and glomeruli.

### U-ExM of human clinical nephrectomy samples

Finally, it was important to determine if our modified version of U-ExM can be applied to expand human clinical kidney specimens for downstream analysis. Healthy and diseased renal tissue sections were obtained from the Kidney Translational Research Center at Washington University in St Louis. We tested samples that were fixed with PFA and/or formalin, since most clinical samples are preserved in this way. We avoided fresh frozen (unfixed prior to cryopreservation) specimens since we observed artifacts associated with expanding these mouse kidney sections (described above). In proof-of-principle studies, we applied U-ExM to healthy human from nephrectomies and from patients with Autosomal Dominant Polycystic Kidney Disease (ADPKD). ADPKD is a well-established ciliopathy that is characterized by defects in a) ciliary structure and function, b) changes in epithelial cell growth, polarity, and cytoskeletal organization, c) the development of large fluid filled cysts that destroys the renal parenchyma, and d) fibrosis and extracellular matrix (ECM) deposition that leads to renal failure (Bastos & Onuchic, 2011; Fragiadaki et al., 2020; Grantham et al., 2011; Hou et al., 2002; Lee & Somlo, 2014; Ma et al., 2017; Norman, 2011; Song et al., 2017; Wilson, 2004; Wilson et al., 1992; Yoder et al., 2002).

We sought to determine whether we could visualize these molecular, organellar, and cellular defects with U-ExM. Using the same antibodies (Table 6), we labeled the centrosome-cilium complexes with acetylated α-tubulin and tubular epithelial cells with E-cadherin, in control and ADPKD samples (Figure 7A-B). As expected, cilia in tubular cysts of ADPKD tissue were longer when compared to cilia in nephron tubules of healthy kidneys (Figure 7C-D) (Huang & Lipschutz, 2014; Shao et al., 2020; Yanda et al., 2023). We also noted the presence of excess centrioles in cystic epithelial cells (Figure 7E), as has been previously described in ADPKD (Battini et al., 2008; Burtey et al., 2008; Cheng et al., 2024; Dionne et al., 2018). There was a pronounced increase in acetylation of the cytoplasmic microtubules in tubular epithelial cells, a posttranslational modification that has been identified in ADPKD cystic kidneys (Figure 7F) (Berbari et al., 2013; Zhou et al., 2014). In addition, there was an increase in ECM deposition and collagen I-positive myofibroblast staining in the ADPKD tissue (Figure 7G) (Fragiadaki et al., 2020; Norman, 2011; Song et al., 2017). In sum, our results demonstrate that U-ExM can be a valuable tool for analysis of cytoskeletal (and other morphological defects) in ADPKD and other human renal diseases.

**Figure 7.**
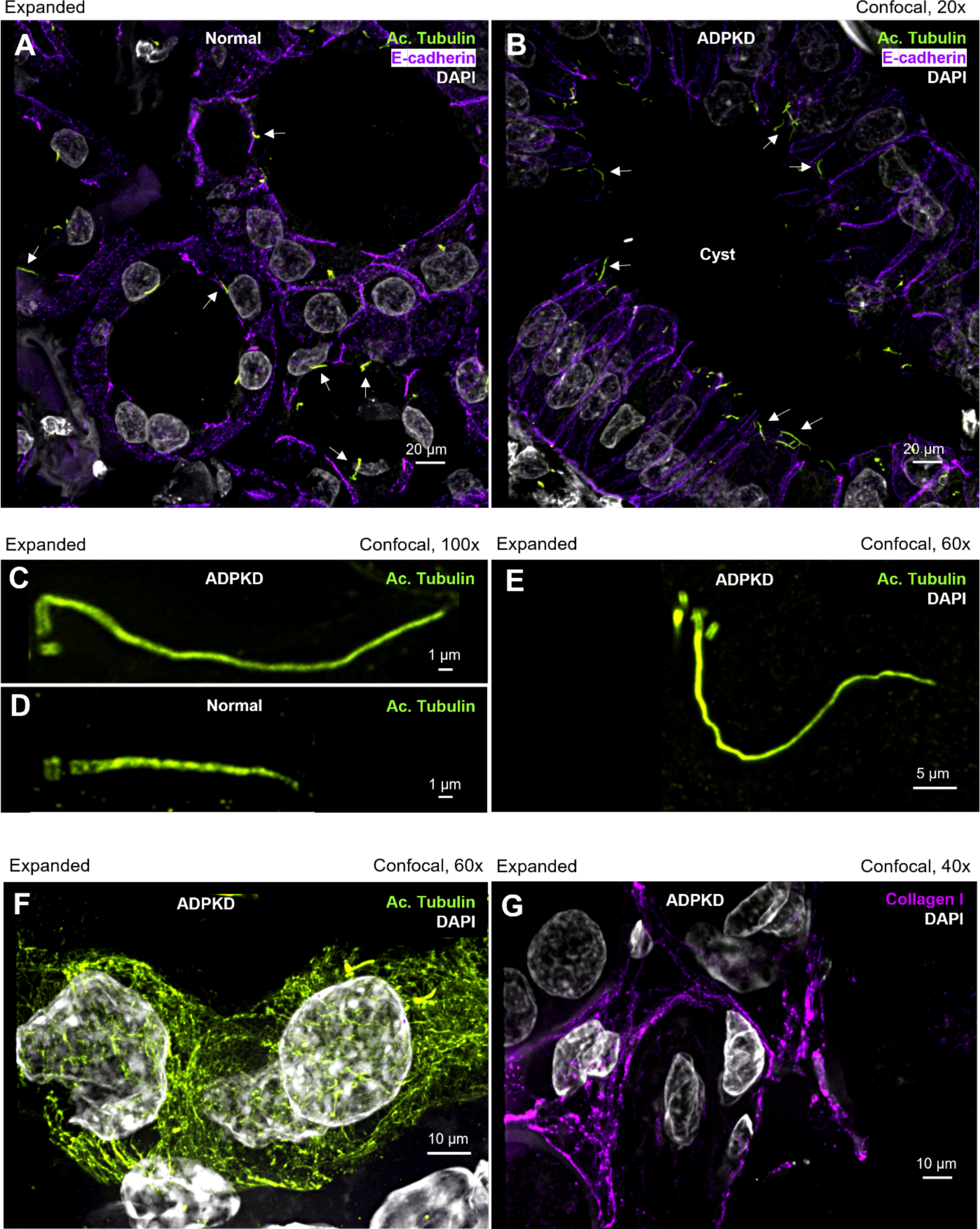
U-ExM of human Autosomal Dominant Polycystic Kidney Disease sections. **(A-B)** Representative immunofluorescence images of normal human kidney (A) and ADPKD sections (B). Centriolar and ciliary microtubules were labeled with antibodies against acetylated α-tubulin (green), nephron tubular epithelia with E-cadherin (purple), and nuclei with DAPI (white). White arrows point to cilia in the normal and cystic tubules. **(C-D)** Example of changes in cilium length in cystic (ADPKD) and normal epithelial cells. **(E)** Centrosome amplification is easily visualized in human ADPKD cystic cells. Centrioles were labeled with acetylated α-tubulin (green). **(F)** Increased acetylation of cytoplasmic α-tubulin in tubular cells in ADPKD. **(G)** Extracellular matrix deposition can be visualized in expanded ADPKD samples using antibodies against Collagen I (magenta).

## Discussion

In this study, we have adapted and modified the U-ExM protocol to expand mouse and human kidney sections. We describe a step-by-step method for expansion, detection and analysis of centrioles, cilia, several cytoskeletal structures and renal cell types. This method achieves ∼4x isotropic expansion and takes 4 days to complete from start to finish. The protocol is straightforward, accessible, and affordable with no need for advanced equipment or microscopes. It allows visualization and analysis of nanoscale-resolution structures like centrioles and cilia, as well as the precise localization of diverse proteins associated with these organelles. Furthermore, it can facilitate the staining and analysis of the extracellular matrix, basement membrane, podocyte slit diaphragm, and several other cytoskeletal elements. Finally, this method can be widely applicable as it enables use with both pre-clinical and clinical tissues preserved in common fixatives, like paraformaldehyde or formalin.

Several approaches to expand and immunolabel kidneys have been developed using the ExM, MAP, ExPath and Magnify methods (Bucur & Zhao, 2018; Chozinski et al., 2018; Klimas et al., 2023; Ku et al., 2016; Kylies et al., 2023; Siegerist et al., 2022; Unnersjö-Jess et al., 2021; Unnersjö-Jess et al., 2018; Wunderlich et al., 2021; Zhao et al., 2017). However, there are several advantages to the U-ExM approach: a) First, U-ExM allows for post-expansion staining of the sample. This is especially important for rare clinical specimens, where the researcher might only have access to a limited number of sections. In the previous approaches (ExM and ExPath), the kidney sections were first immunostained with antibodies then expanded using the protease digestion method. The major disadvantage is that only one combination of staining can be used before the expansion step. In contrast, U-ExM will provide multiple gel punches from a single kidney section, which can easily facilitate 10-15 staining combinations; b) Another advantage of U-ExM is the use of less antibodies. ExM and ExPath require high density immunolabeling, which might not be possible for some proteins due to insufficient epitope accessibility, or the need for high volumes of costly (and potentially rare) antibodies to be used. In contrast for U-ExM, staining can be performed on small pieces of gel, thus decreasing the need for large amounts of antibodies; c) U-ExM involves tissue staining post-denaturation, which can improve detection of some epitopes due to enhanced accessibility; d) U-ExM allows for imaging of the ‘real’ immunolabeled protein in the tissue, compared to previous methods where only the image of the expanded fluorophore imprint (after total or partial protein digestion) is acquired; and e) We found that approximately 80% of antibodies tested in our laboratory worked well and specifically labeled the associated structures in expanded tissue, highlighting the broad utility of the approach. Finally, U-ExM is advantageous to electron microscopy for analyses of kidney tissues since it allows visualization of both morphology and protein composition. Protein localization by EM traditionally requires immunogold labeling, and it is difficult to define the positions of more than one protein at a time. Moreover, EM requires specialized expertise and equipment, which may not be accessible to most researchers.

In conclusion, this protocol presents a simple step-by-step method for renal tissue expansion that can be applied to both pre-clinical and clinical samples, allowing comprehensive structural analysis of cytoskeletal structures in normal and diseased kidney tissues.

## Authorship

E.L. performed most of the experimental work, data acquisition and quantitation, prepared the figures, and wrote the manuscript. P.P. and D.Y. performed experimental work on glomeruli, analyzed data, and prepared figures. R.P. and J.W. provided technical expertise with hydrogel expansion experiments, and shared reagents. H.S., J.M., A.H., S.D. and S.B. provided reagents, technical support, and critical feedback. M.R.M. supervised all aspects of the project and helped write the manuscript. All authors read and edited the manuscript.

## Acknowledgments

We would like to thank Dr. Sanjay Jain at the Kidney Translational Research Center (Washington University in St Louis) for providing the de-identified ADPKD kidney sections. We also thank members of the Mahjoub lab and the Washington University Ciliopathy Research Group for helpful advice and feedback on this project, as well as critical reading of the manuscript. Some figure panels were created with BioRender.com. We acknowledge the Washington University Center for Cellular Imaging (WUCCI) for assistance with some image acquisition, supported in part by the Washington University School of Medicine, the Children’s Discovery Institute and St. Louis Children’s Hospital (CDI-CORE-2015-505 and CDI-CORE-2019-813), and the Foundation for Barnes-Jewish Hospital (3770 and 4642). This study was supported by funding from the Department of Defense PRMRP (W81XWH-20-1-0198) to MRM., the National Institute of Diabetes and Digestive and Kidney (NIH DK131177) for JHM and HYS, and the National Heart, Lung and Blood Institutes (NIH HL128370) to SLB, SKD and MRM.

## Conflict of Interest statement

The authors declare no competing financial interests.

## Notes

### Competing Interest Statement

The authors have declared no competing interest.

